# Development of bioactive electrospun scaffolds suitable to support skin fibroblasts and release *Lucilia sericata* maggot excretion/secretion

**DOI:** 10.1101/864892

**Authors:** Annesi G Giacaman, Ioanna D. Styliari, Vincenzo Taresco, David Pritchard, Cameron Alexander, Felicity R Rose

## Abstract

Larval therapy has been reported to exert beneficial actions upon chronic wound healing by promoting granulation tissue formation, antimicrobial activity and degrading necrotic tissue. However, the use of live maggots is problematic for patient acceptance, and thus there is a need to develop materials which can adsorb and release therapeutic biomolecules from maggot secretions. Here we describe the fabrication of a novel bioactive scaffold that can be loaded with *Lucilia sericata* maggot excretion/secretion (*L. sericata* maggot E/S) for wound therapy, and which also provides structural stability for mammalian cell-growth and migration. We show that electrospun scaffolds can be prepared from polycaprolactone-poly (ethylene glycol)–block copolymer (PCL-b-PEG) blended with PCL, to form fibres with average diameters of ~4 μm. We further demonstrate that the fibres are able to be loaded with *L. sericata* maggot E/S, in order to influence fibroblast migration through protease activity. Finally, we show that after 21 days, the cumulative amount of released *L. sericata* maggot E/S was ~14 μg/mL from PCL-b-PEG/PCL scaffolds and that the protease activity of *L. sericata* maggot E/S was preserved when PCL-b-PEG/PCL scaffolds were used as the release platform.

## Introduction

Larval therapy has been reported to exert beneficial actions upon chronic wound healing [1, 2]. The alimentary fluid from *Lucilia sericata* excretion/secretion (*L. sericata* maggot E/S) contains a range of serine proteinases, which can degrade necrotic tissue [3], disinfect the wound sites [4] and promote granulation tissue formation at the wound site [5]. The beneficial effect of larval excretion/secretion on wound healing is strongly supported by 8 clinical trials [6–8], supported by *in vitro* studies [9]. The FDA (Food and Drug Administration, USA) approved *L. sericata* maggot E/S therapy as a medical device in January 2004 [10]. In the same year, the National Health Service in the UK approved the prescription of sterile *L. sericata* maggot for debridement therapy [7]. However, there are many limitations of the use of sterile *L. sericata* maggots for wound healing. These include i) maggots require a moist environment to survive, for this reason the use of *L. sericata* maggots is accepted exclusively for treatment of exudative wounds and contraindicated on dry wounds ii) maggots produce malodour and discomfort to the patient and iii) even though they are only used when located within a porous bag, there is poor acceptance by patients and by medical staff [11]. For this reason, research groups have tried to incorporate the active molecules from *L. sericata* maggot E/S in pharmaceutical formulations. Prototype hydrogels containing recombinant *L. sericata* chymotrypsin have been developed and were shown to accelerate wound closure *ex vivo* [12]. In addition, *L. sericata maggot* E/S has been incorporated and released in a PVA hydrogel. Findings showed a significant enhancement on the rate of wound closure (*in vitro* scratch assay) when the wounds were exposed to *L. sericata* maggot E/S released from the hydrogel compared with control [13]. However, hydrogel formulations have 90 % water content and are their used can result in skin maceration and bacterial colonization if wounds for treatment are highly exudative and this in turn can delay/impair the wound closure [14].

Accordingly, there is a need for more robust, non-hydrogel, formulations *L. sericata* maggot E/S which could be used to support fibroblast proliferation and migration in accelerated wound healing applications.

Electrospun scaffolds provide a potential means to meet this need, as they have solid, fibrous and porous structures that can serve as a substitute for the native extracellular matrix [4]. Electrospinning is a versatile technique that allows the fabrication of nano- and micro-fibres by modifying the process parameters [5]. Many synthetic polymers have been used to generate fibrous scaffolds though electrospinning of which the most widely-applied in medical applications are those on poly(lactic acid) (PLA) and polycaprolactone (PCL), either as homopolymers or as blends with poly (ethylene glycol) (PEG) [6]. This is because PLA and PCL have been authorised for clinical application in drug delivery devices [7], while Poly (ethylene glycol) (PEG) is component of many existing medical and consumer products. These synthetic polymers are advantageous for clinical purposes as they are relatively cheap compared to highly purified natural polymers, and synthetic materials can be more easily tuned to enhance specific properties such as mechanical strength, shape and biodegradation rate [8]. Fibrous scaffolds derived from these polymers can be further enhanced to be bioactive through the incorporation of therapeutic agents which can then be released as the fibres swell or degrade in biological media. Bioactive fibrous scaffolds refer to a matrix or scaffold that provide structural support for cells and with the intention of guiding the cellular response by delivering therapeutic molecules [10]. Bioactive electrospun scaffolds have been used in areas such as cancer treatment for the release of doxorubicin, and in cardiovascular disease to release bioactive agents (i.e growth factors, heparin, anti-inflammatories) [11]. In wound healing, the most common loaded drugs are antibiotic [12], herbal extracts [13] and proteins (i.e growth factors) [14]. Furthermore, while some synthetic polymers can present challenges with respect to biocompatibility due to the lack of specific cell-recognition motifs and sub-optimal cell-surface interfacial interactions, [9] they can be surface-modified post-spinning with appropriate functionalities if required to encourage cell attachment and growth [15].

Here, we describe an approach to bioactive scaffolds designed to allow the sustainable release of *L. sericata* maggot E/S from the electrospun fibres as well as support for cell proliferation and migration. polycaprolactone-poly (ethylene glycol)–block copolymer (PCL-b-PEG) blended with PCL (PCL-PEG/PCL) was electrospun to create the fibrous scaffold, with PEG included within the scaffolds to enhance the wettability and cytocompatibility of the surface, and to accelerate the degradation rate of the polymer, and thereby improve *L. sericata* maggot E/S release. To test these hypotheses, the *in vitro* biocompatibility and migratory behaviour of 3T3 GFP mouse dermal fibroblasts and human BJ6 fibroblasts, on electrospun PCL and PCL-b-PEG/PCL scaffolds were investigated.

*L. sericata* maggot E/S was incorporated in the scaffolds by post electrospinning modification. Release evaluation was carried out by immersing the loaded-scaffolds in PBS for 21 days. And the relative bioactivity of *L. sericata* maggot E/S was confirmed by examining protease activity after the release.

We believe these data demonstrate that effective biomaterial can be produced by electrospinning, that it can incorporate and release *L. sericata* maggot E/S bioactive molecules, and that the fibrous scaffolds can enhance cellular infiltration and proliferation

## Materials and Methods

### Development of electrospun scaffolds

Electrospinning electrospun scaffolds were fabricated in a commercial electrospinning apparatus EC-CLI (IME Technologies). PCL scaffold: scaffolds were electrospun using PCL (80 KDa; Sigma Aldrich) dissolved in chloroform (8% *w*/*w*) (Life technology, Fisher scientific). PCL scaffolds were used as controls to compare properties with PCL-b-PEG/PCL scaffolds. Monomethoxy poly (ethylene glycol) (mPEG) ^(^5 KDa), ɛ-caprolactone, and 1,5,7 -Triazabicyclo (4.4.0) dec-5-ene (TBD) were purchased in Sigma Aldrich and used to synthesised PCL-b-PEG (57KDa) prepared by ring–opening polymerization. In-house synthesised PCL-b-PEG (57 KDa) was blended with PCL (80 KDa) in a proportion of 40/60 dissolved in chloroform, with a final solution concentration of 15% (*w*/*w*). Polymeric solutions were transferred to a syringe using a capillary tube spinneret of 0.6 mm inner diameter which was arranged vertically. Temperature was set at 20°C and 44% relative humidity. Electrospinning parameters were set as described in complementary data.

### Scaffold characterization

#### Fibre morphology and scaffold thickness

Morphology of the fibres and diameter distribution were evaluated by SEM images, taken with a FEI- XL30 (tungsten filament) scanning electron microscope (Philips). The diameter of individual fibres were measured by using imageJ software (1.48 v, USA National Institutes of Health). A total of five images (at 3000x magnification) from five random fields were used for a total of 100 measurements. For each scaffold, three different samples were taken from top to bottom of the electrospun sheet. Measurement of scaffold thickness were carried out by freezing scaffold samples at −80°C, and after 24 h a cross section of the samples was excised using a cold scalpel blade. Samples were oriented tangentially to the surface of the SEM sample holder and imaged as described above. For fibre diameter analysis, one hundred measurements were taken using imageJ software. Frequency distribution and non-linear regression (Log Gaussian) was analysed using Prism (GraphPad Prism6, San Diego CA).

#### Scaffold Porosity

The porosity of the electrospun scaffolds was assessed using micro computed tomography (micro-CT) (Skyscan 1174, Skyscan, Belgium). The samples (in triplicate from 3 different scaffolds) were set vertically inside of a straw and fixed into the top side with blue glue (Blu-tack). The images were binarized with a threshold range 22 to 255 (gray values). A misalignment of −5 to +5 was used. 3-D measurements and structural analysis were performed with CTAn software (Bruker software, SkyScan 1174). NRecon (Bruker software, Skycan 1174) was used to reconstruct 2D slices. In addition, pore size was evaluated.

#### Water Contact Angle (WCA)

Measurements of the WCA at the surface of the electrospun scaffolds were carried out using a modular CAM 200 optical contact angle meter (KSV Instruments Ltd) equipped with a camara (JAi^®^). Rectangular areas (3×1.5 cm) of the scaffolds PCL and PCL-PEG/PCL (average fibre diameter 4 μm) were fixed to glass cover slips, 3 pieces of the same scaffold were analysed, and measurements repeated 3 times. The sessile drop method, advancing (θa) and receding (θr) analysis were performed (Korhonen et al., 2013).

#### Mechanical properties/uniaxial tensile properties

Mechanical properties of the electrospun scaffolds were determined with a uniaxial testing machine (TA-HD Plus Texture Analyser, stable Micro System Instron 3345). A 5 Kg load cell under a cross-head speed of 10 mm/min at 25°C was used. All scaffolds were cut in a ‘dog bone’ shape with dimensions of the rectangle of 15×10 mm. The thickness were measured with a digital calliper having a precision of 1 μm and confirmed by SEM measurements. At least in triplicate from 3 different scaffolds. Only the scaffolds that had broken in the centre were considered for statistical analysis. Young’s modulus, the yield point, the elongation at break and ultimate tensile strength were evaluate and graphs of share rate vs viscosity and shear rate vs shear stress were generated.

#### *In vitro* degradation studies

Evaluation of scaffold degradation were carried out using 0.8 mm scaffolds of known weight and immersing them in 5 mL of Phosphate buffer saline (PBS) during agitation at 37°C. Scaffolds were reweighed regularly (0, 1,2,3 and then 5,7,9,11 and 12 months). Additionally, selected samples were taken at time 0, 5 months and 12 months for Gel permeation chromatography (GPC) analysis. One individual experiment in triplicate was carried out.

### Biological evaluation of the scaffolds

#### Mammalian cell maintenance

NIH 3T3, GFP -3T3 mouse embryo fibroblast cells and BJ6 foreskin human fibroblasts cells (passage number 10) were provided by Dr. James Dixon (The University of Nottingham) and were previously purchased from ECACC. Fibroblast were cultured in Dulbecco’s modified Eagle’s medium (DMEM) supplemented with foetal calf serum concentration 10 %, *v*/*v* and antibiotics /antimitotic final concentration (100 units/mL penicillin G, 100 μg/mL streptomycin sulphate and 0.25 μg/mL amphotericin (all from Sigma Aldrich) at 37°C, 5% CO_2_ in air at 95% relative humidity. Sub confluent cells were harvested with trypsin-EDTA and used for further experiments. Fresh medium was replenished every second day. For washing procedures was used Hank’s balance salt solutions (HBSS) (Gibco/Invitrogen Life Technologies).

#### Scaffold preparation and cell seeding

To prepare scaffolds for mammalian cell culture, circles of PCL and PCL-PEG/PCL scaffolds, with a diameter of 8 mm were cut from 3 different random areas using a metallic puncher. The scaffolds were fixed to the bottom of a 24 non-tissue culture treated plates with a small quantity of aquarium seal glue. Later, scaffolds were sterilised using a UV lamp (CL-100UV) for 10 min at power 200 (x100 μJ/cm^2^). Then, scaffolds were immersed in 1 mL of cell culture media and maintained at 37°C, 5% CO_2_ in air at 95% relative humidity for 24 h. After that, the cell culture media was replaced and scaffolds were seeded with 60,000 cells/well, maintained at 37°C, 5% CO_2_ in air at 95% relative humidity.

#### Cell viability

The metabolic activity of NIH 3T3 GFP and BJ6 fibroblasts on each scaffold was determined by performing the AlamarBlue™ assay (Life technologies, Fisher scientific) which contains a fluorometric/colorimetric indicator (resazurin) that undergoes into florescence and colour changes (resorufin) in response to chemical cellular metabolic reductions. The AlamarBlue™ assay was carried out after 6 h, 2 days, 4 days, 6 days, 9, 12 days and 15 days post seeding. Scaffolds were transferred to new non-tissue culture treated plate before carrying out the AlamarBlue™ assay to avoid measuring cells attached to the bottom of the well plate. Relative fluorescence units were measured in a plate reader (Tecan Infinite M200, Reading), set at an excitation wavelength of 560 nm and emission at 590 nm. Experiments were carried out in triplicate for a minimum of n=3 individual experiments.

#### Cell infiltration into the Scaffolds

Evaluation of the extent of NIH 3T3 GFP infiltration into the scaffolds after 5 days of culture was conducted by z-stacked confocal microscopy (LSCM; Zeiss, LSM880) analysis (pinhole set to 1 Airy unit, pixel dwell time: 0.26 μs, 5 μm plane thickness). Laser excitation at 405 nm and 488 nm were used, and sequential imaging of Hoechst (blue) and GFP (green fluorescent protein) was performed. Experiments were carried out in triplicate from at least three different scaffolds The infiltration depth (%) of the fibroblasts was calculated by estimating the maximum distance (microns) migrated by fibroblasts from the top to the bottom of the scaffolds (visualization on Z-stack) and using the total distance, the thickness of the scaffolds estimated by SEM [16].

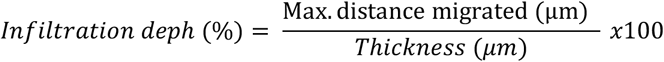

### PCL-b-PEG/PCL scaffolds as a platform for protein release

#### Protein loading efficiency studies

*L. sericata* maggot E/S were collected from sterile, freshly hatched *L. sericata* maggot (LarvE, Surgical Materials Testing Laboratory, Cardiff, UK) as reported by Horobin and colleagues [17].

*L. sericata* maggot E/S was labelled with NHS-Rhodamine (thermo scientific Cat. Number 46406) and the study of adsorption of protein onto the scaffolds carried out. NHS-rhodamine (7 mg) was dissolved in 1 mL dimethyl sulfoxide (DMSO) (Scientific laboratory supplier) and stirred for1 h, in the dark, at 25 °C. The solution was added under agitation to the *L. sericata* maggot E/S solution for 1 h, in the dark at 25 °C. Unreactive NHS-rhodamine was removed by dialysis using a membrane with 3.5 K MWCO (Snakeskin ™ dialysis tubing) against deionized water. Dialysis membranes were embedded in 3 L of deionised water, which was changed 3 times for 4 days, in the dark at 25°C

Scaffolds with similar weight (800 μg) were cut into circles and prepared as described above. The initial concentration of absorbed protein labelled with rhodamine was quantified by fluorescence spectroscopy analysis and confirmed by the Bradford assay according to manufacturer’s instructions (Biorad). Scaffolds were fully immersed in 1 mL of protein solution and incubated at 37°C with constant agitation for 24 h. The amount of protein absorbed on the scaffolds was determined indirectly by measuring the protein concentration in the supernatant after 24 h. The following equation was used to calculate the absorption efficiency [18]. Experiments were carried out in triplicate for a minimum of n=3 individual experiments.

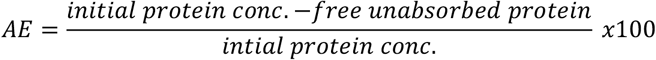

#### Release of proteins

The release profile of *L. sericata* maggot *E/S* from PCL and PCL-b-PEG/PCL scaffolds was assessed by washing scaffolds 3 times with deionised water to remove excess protein and dried in a vacuum oven at 25°C for 48 h. For the release studies, protein loaded-scaffolds were immersed in 1 mL PBS (pH 7.4). The samples were maintained under agitation at 37°C 5% CO_2_ in air for 21 days. At each time point (30 min, 1-6 h hourly,12 h, day 1-6 daily, every 3 days until day 21) 1 mL was removed and replaced with fresh; the samples were stored at −20°C for further protein quantification and activity estimation. Experiments were carried out in duplicate for a minimum of n=3 individual experiments. Protein release was expressed as cumulative release % as a function of time and calculated using the following equation [19]:

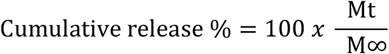

Where Mt is the amount of protein released at time t and M∞ is the total amount of protein adsorbed onto the scaffold.

#### Quantification of proteins

The concentration of protein absorbed and released from the scaffolds were performed by fluorescence spectroscopy and Bradford Assay. The *L. sericata* maggot E/S-NHS rhodamine conjugate was measured at Ex/Em 552/575 nm and plotted as relative fluorescence units (RFU). For Bradford assay was used two different absorbance 590- and 450-mm. Experiments were carried out in triplicate for a minimum of n=3 individual experiments.

#### Relative bioactivity of *L. sericata* maggot E/S

The bioactivity of *L. sericata* maggot E/S after release was evaluated by pooling all the aliquots collected at the different time points. The samples were concentrated using an ultrafiltration device (Vivaspin) with a MWCO membrane of 3 KDa and centrifugation at 3000 xg for 30 min at 4°C.

The bioactivity of the *L. sericata* maggot E/S was assessed using a Protease Fluorescent Detection Kit (Sigma Aldrich), This kit detects the activity of serine, metallo and aspartic protease activity by using a FITC-casein substrate. The action of the protease is the cleavage of the casein-FITC bond releasing free FITC which can be detected [20]. Experiments were carried out in triplicate for a minimum of n=3 individual experiments.

#### Statistical analysis

All data was expressed as mean±SD. Experiments were carried out in triplicate for a minimum of n=3 individual experiments, unless otherwise indicated. To analyse statistically significant differences with respect to cell viability between different scaffolds, protein loading and release from the scaffolds, a Two way-ANOVA with Turkey’s multiple comparison post-hoc test was performed using Prism (GraphPad Prism6).

## Results

### Improved cellular infiltration into scaffolds with PEG content and fibre diameter in a microscale range

PCL alone and PCL-b-PEG/PCL electrospun fibres were fabricated with an average fibre diameter of ~4 μm. Figure 1 shows representative SEM images of the PCL-b-PEG/PCL and PCL microfibre scaffolds and image analysis determined the average fibre diameter to be 3.5 ±0.4 and 3.6±0.8 μm respectively.

**Figure 1.**
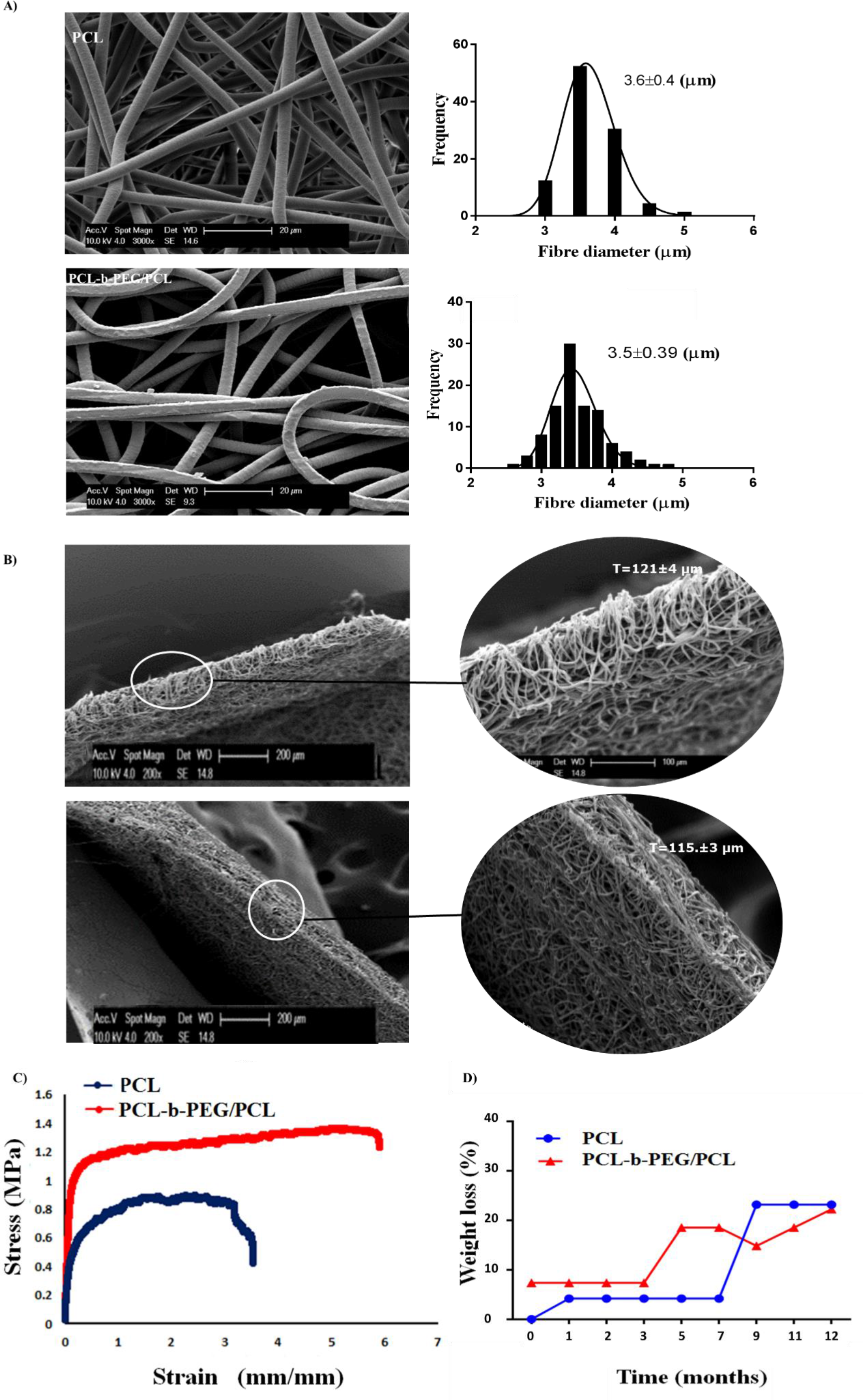
**A)** Representative SEM micrographs (at x3000) of PCL-PEG/PCL and PCL scaffolds produced under different conditions (included in supplemental data). Scale bar20 μm and log -normal distribution of the size and frequency of their respective fibres. **B)** PCL and PCL-PEG/PCL scaffold thickness. Representative scanning electron micrographs of cross-sections of PCL and PCL-PEG/PCL electrospun scaffolds (at x 200 and x100 magnification). **C)** Representative stress-strain curves of PCL and PCL-b-PEG/PCL scaffolds(no error bars), **D)** Degradation rate study-graph of weight loss of PCL and PCL-b-PEG/PCL scaffolds over a period of 12 months(no error bars), one independent experiment carried out in triplicate. Refer to table 2 for GPC analysis of degradation rate.

PCL scaffolds were produced with a thickness of 121±4 μm and a porosity of 58±2 %, and PCL-b-PEG/PCL scaffolds were similar at 115±3 μm thickness and 67±3 % porosity.

It was hypothesised that the scaffold composed of PCL-b-PEG/PCL fibres may exhibit enhanced wettability and thus enhanced degradation rate and cell-compatibility. Table 1 shows a summary of the scaffold parameters including the static contact angle, advancing and receding water contact angle for scaffolds PCL and PCL-b-PEG/PCL. Contact angle measurements confirmed that only the PCL scaffolds are hydrophobic (108°) whilst the PCL-b-PEG/PCL scaffolds exhibited a water contact angle of 100° with no statistical significances found. (Table 1).

**Table 1.** A summary of the main properties of PCL-b-PEG/PCL and PCL scaffolds including porosity, advancing/ receding contact angle, hysteresis and static contact angle, mechanical properties and degradation. Experiments were carried out in triplicate and data were express as mean±SD.

| Material | Fibre diameter<br>µm | Porosity<br>% | Contact angle ° |  | Mechanical properties |  |  |  | Degradation of scaffolds |  |  |  |  |  |
| --- | --- | --- | --- | --- | --- | --- | --- | --- | --- | --- | --- | --- | --- | --- |
|  |  |  |  |  |  |  |  |  | Mw |  | Polydispersity<br>Mw/Mn |  | Polydispersity<br>Mz/Mn |  |
|  |  |  | Static<br>contact<br>angle, | Advancing<br>contact<br>angle, °<br>Θα | Yield point<br>(MPa) | Young's<br>modulus<br>(MPa) | Elongation<br>at break<br>(mm/mm) | Ultimate<br>tensile<br>strength<br>(MPa) | Initial | After 5<br>months | Initial | After 5<br>months | Initial | After 5<br>months |
| PCL | 3.6±0.4 | 58±2 % | 108±5 | 108±3 | 0.6±0.1 | 6.0±0.8 | 3.4±0.2 | 0.9±0.4 | 2.9x10<sup>5</sup> | 2.5x10<sup>5</sup> | 1.2 | 1.4 | 1.4 | 1.9 |
| PCL-b-PEG/PCL | 3.5±0.4 | 67±3 % | 101±3 | 99±29 | 1.0±0.1 | 8.8±0.8 | 5.1±0.9 | 0.9±0.5 | 2.5x10<sup>5</sup> | 1.6x10<sup>5</sup> | 1.3 | 2.4 | 1.6 | 4.0 |

Figure 1 shows stress-strain curves of PCL and PCL-b-PEG/PCL electrospun scaffolds, using uniaxial tensile tests. In general, PCL and PCL-b-PEG/ PCL scaffolds presented a similar stress-strain behaviour which indicated that the scaffolds were flexible and deformable. Young’s modulus, yield point, and elongation at break are summarized in Table 1. The elongation at the break point was 5.1±0.9 mm/mm for PCL-b-PEG/PCL and 3.4±0.2 mm/mm for PCL scaffolds. Moreover, in terms of Young’s modulus, PCL-b-PEG/PCL scaffold reached values of 8.8±0.8 MPa, compared with PCL electrospun scaffolds, with average values of 6.0±0.8 MPa which was statistically significantly different. The ultimate strength of PCL-b-PEG/PCL and PCL scaffolds were similar reaching values of ~0.9 MPa.

*In vitro* degradation studies were carried out with the PCL scaffold losing 5% of its weight in the first month. After that, the weight was constant for 7 months with a further weight loss of around 3% and again the weight was maintained. These results were in agreement with GPC analysis of molecular weight and polydispersity which indicated that both scaffolds experienced degradation. (Table 1). PCL slightly decreased in molecular weight from 2.9×10^5^ to 2.5×10^5^ Mw and increased in terms of polydispersity Mw/Mn from 1.2 to 1.4 over 5 months. PCL-b-PEG/PCL scaffolds showed a greater weight loss (around 10%) within the first month. After that, the electrospun scaffolds lost around 20% of their weight in of the following 9-month period. Complementary GPC analysis showed a decrease in molecular weight for the PCL-b-PEG/PCL scaffold from 2.5×10^5^ to 1.6×10^5^ Mw with an increase in polydispersity Mw/Mn from 1.3 to 2.4 over a period of 5 months.

The 3T3 fibroblasts cultured on the PCL-b-PEG/PCL scaffolds proliferated over a 12-day period and showed a significantly higher metabolic activity (p <0.0001) than for those cultured on PCL scaffolds on day 12. No statistically significant differences were found in terms of metabolic activity between fibroblasts cultured on PCL-b-PEG/PCL scaffolds and PCL scaffolds during the first 9 days of culture (Figure 2-A). Figure 2-B compares the metabolic activity of human BJ6 fibroblasts on PCL and PCL-b-PEG/PCL scaffolds over 15 days of culture. BJ6 fibroblasts cultured on PCL scaffolds did not exhibit an increased metabolic activity over 15 days of culture suggesting that they did not proliferate. Metabolic activity on the PCL-b-PEG/PCL scaffolds was significantly higher, illustrating the ability of BJ6 cells to proliferate compared with those cultured on PCL.

**Figure 2.**
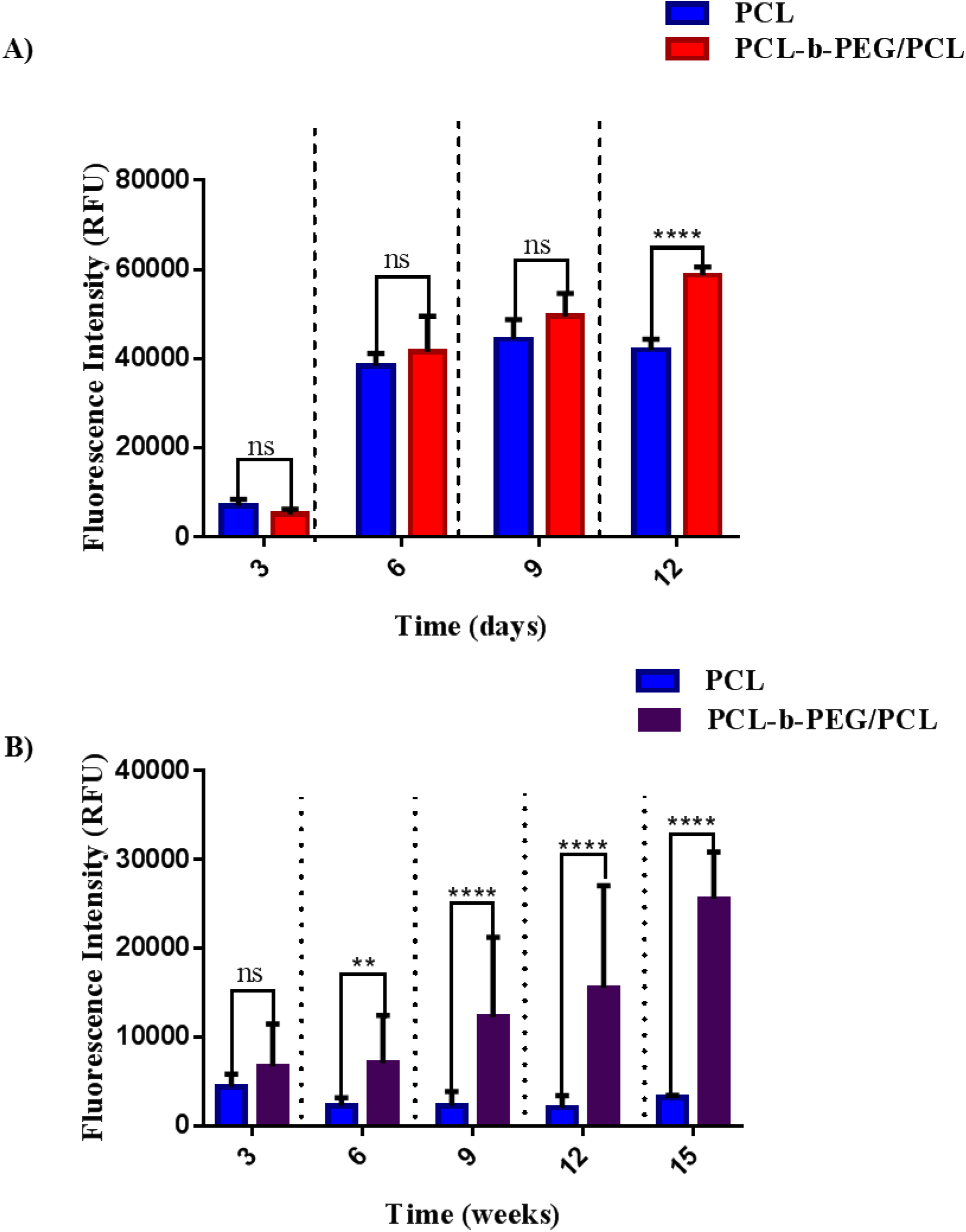
Fibroblast metabolic activity when cultured on the scaffolds was assessed using AlamarBlue™. **A**) Fluorescence readings of metabolic activity of 3T3 GFP fibroblasts cultured on PCL and PCL-b-PEG/PCL scaffolds, over 12 days. **B**) Fluorescence readings of metabolic activity of dermal human fibroblasts BJ6 cultured on PCL and PCL-b-PEG/PCL over 15 days of culture. Two way-ANOVA, Tukey’s multiple comparison test. Three independent experiments carried out in triplicate (means±SD). Ns for P>0.05, * for P≤0.05, ** P≤0.01, *** P≤0.001, **** P≤0.0001.

Figure 3 shows the infiltration of cells cultured on the PCL and PLC-b-PEG/PCL scaffolds, which compares the percentage of fibroblast infiltration with respect to the total thickness of the scaffolds. Fibroblast infiltration on the PCL-b-PEG/PCL scaffold was significantly higher (p≤0.0001) (60%) compared with fibroblast infiltration on the PCL scaffold (40%) (Figure 3-B). In summary, the highest fibroblast infiltration and highest metabolic activity was seen when cells were cultured on the PCL-b-PEG/PCL scaffolds in comparison to scaffolds fabricated from PCL alone.

**Figure 3.**
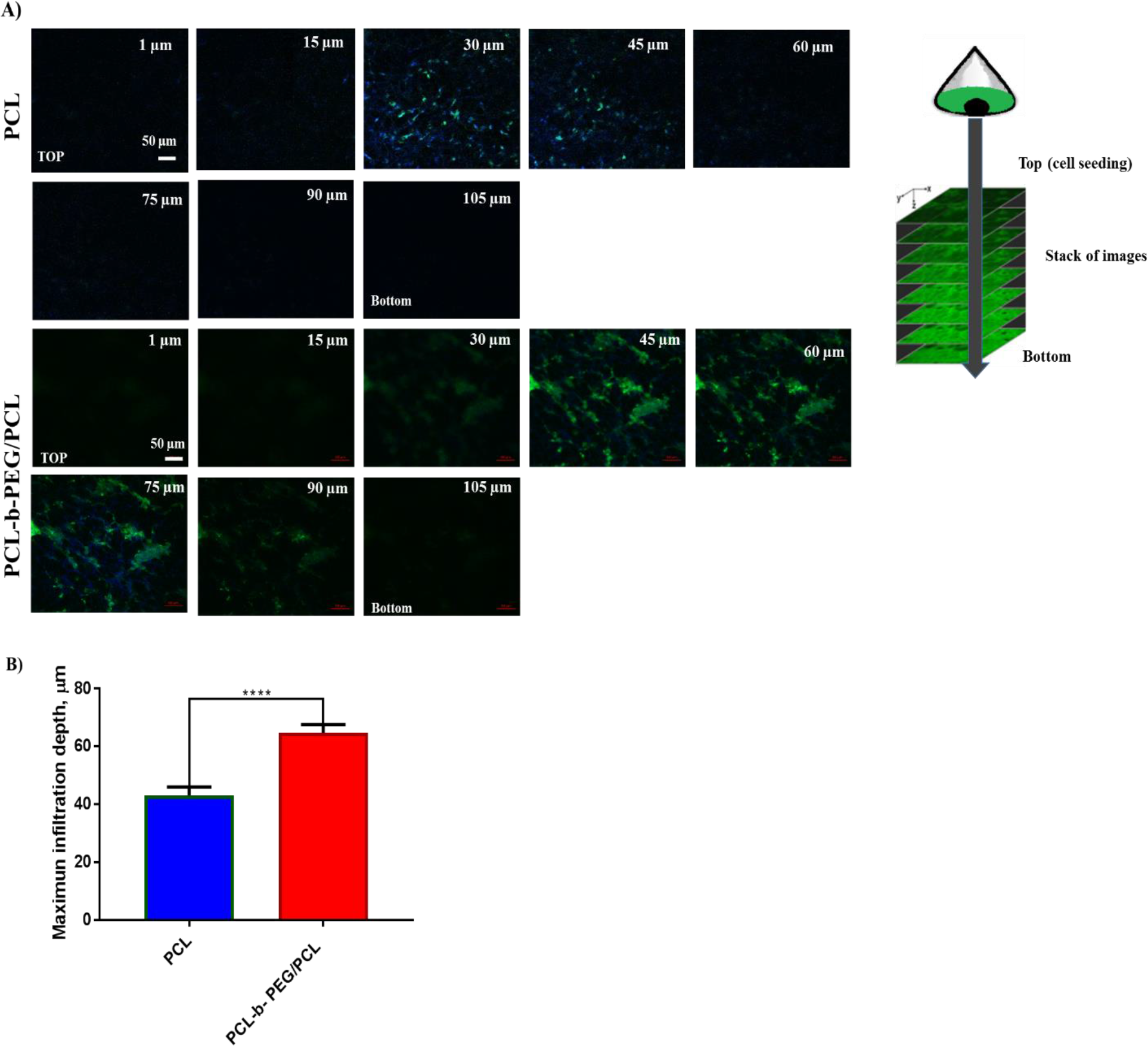
**A)** Confocal microscopy z-stack images of 3T3 GFP fibroblasts seeded onto the PCL and PCL-b-PEG/PCL microfibrous scaffolds after 5 days of growth. In each group, z-stack images at different planes from the cell seeded surface to the bottom are shown with 5 μm plane thickness. Cells expressed GFP (green) and nuclei were stained with Hoechst (blue). Magnification x20, scale bar = 50 μm. **B)** Infiltration depth expressed as a percentage of the total scaffold thickness of 3T3 GFP 3T3 fibroblasts cultured on PCL and PCL-b-PEG/PCL scaffolds, with the same average fibre diameter, over 5 days. Experiments were carried out in triplicate from at least three different scaffolds (means±SD).

### Scaffolds as a platform for the release of *L. sericata* maggot E/S

Figure 4 shows fluorescence images of *L. sericata* maggot E/S-labelled with NHS rhodamine adsorbed onto PCL and PCL-b-PEG/PCL scaffolds following different periods of exposure. Figure 4-B shows a quantitative analysis of *L. sericata* maggot E/S loaded on PCL and PCL-b-PEG/PCL scaffolds. An initial concentration of maggot ES was determined, and it was 40 μg/mL. No significant differences were found in terms of percentage of *L. sericata* maggot E/S loaded on PCL-b-PEG/PCL scaffolds (53.5%) compared with PCL scaffolds (51.9%).

**Figure 4.**
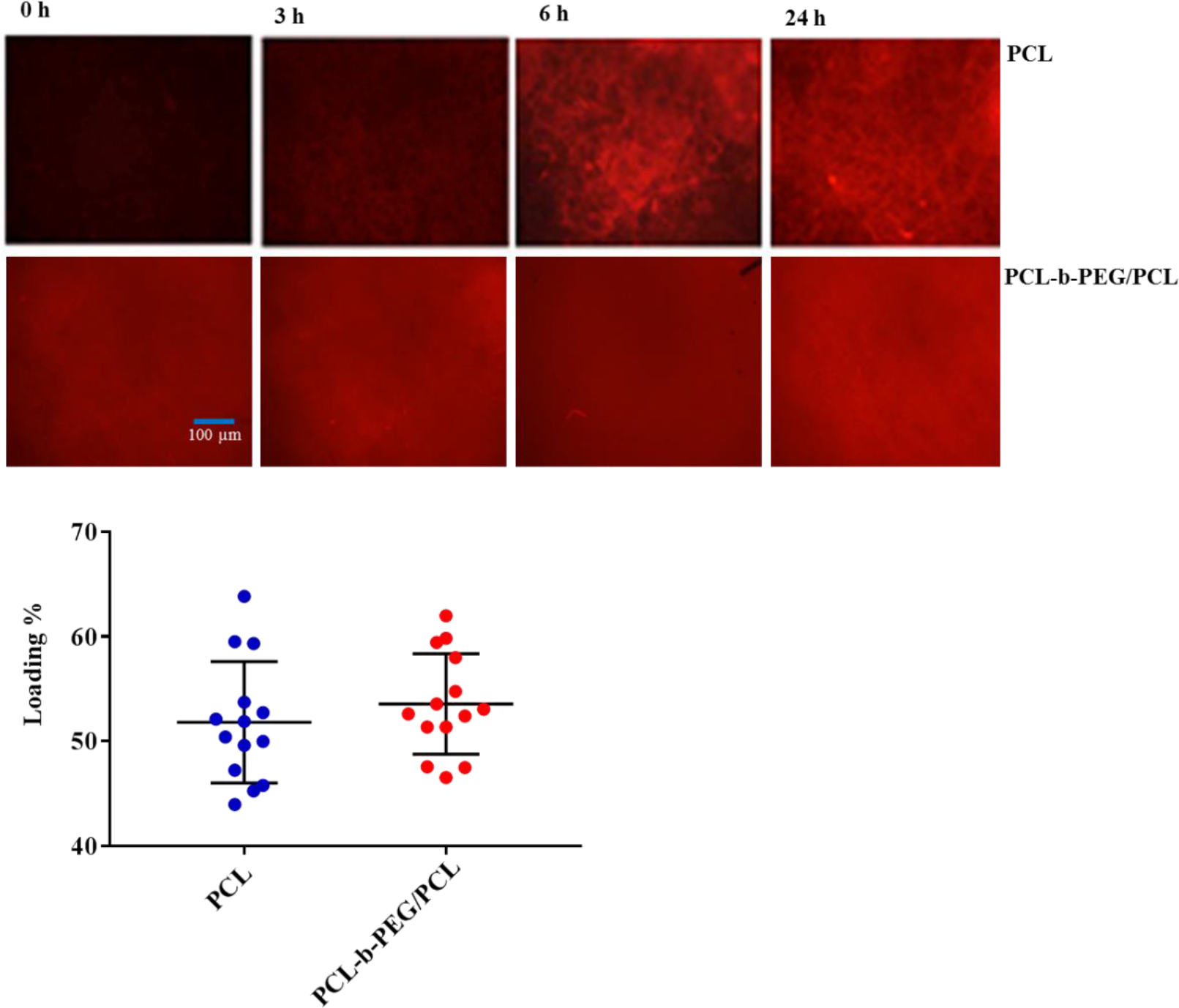
Representative fluorescence microscopy images of *L. sericata* maggot-NHS rhodamine conjugate protein absorbed onto PCL and PCL-b-PEG/PCL scaffolds following different periods of exposure. Magnification 10x, scale bar 100 microns. B) *L. sericata* maggot E/S loading on PCL and PCL-b-PEG/PCL scaffolds. Three independent experiments carried out in triplicate (means±SD).

Figure 5-B shows *L. sericata* maggot E/S release studies from the scaffolds over a 21 days time course. The release of *L. sericata* maggot E/S from PCL began immediately after the immersion of the scaffold in the media (PBS), reaching values of cumulative release of 3.5% of the original loading. After 6 h, *L. sericata* maggot E/S reached values of accumulative release of 25% from PCL-b-PEG/PCL scaffolds, which was slightly higher compared with 22% from the PCL scaffold (P 0.0027). By 24 h, the release of *L. sericata* maggot E/S from PCL-b-PEG/PCL (32%) was slightly higher (P <0.0001) than that released from PCL (28%). *L. sericata* maggot E/S release increased progressively for 21 days, reaching values of 58% from (PCL) and 67% from PCL-b-PEG/PCL scaffolds. PCL-b-PEG/PCL released a statistically significantly higher percentage of *L. sericata* maggot E/S after 21 days compared with that released from PCL (P <0.0001). In terms of cumulative amount of protein released, ~12 μg of *L. sericata* maggot E/S were measured from both electrospun scaffolds after 24 h. After 21 days of release, a further 12 μg of protein was released in total from the PCL scaffolds and 14 μg from PCL-b-PEG/PCL scaffolds.

**Figure 5.**
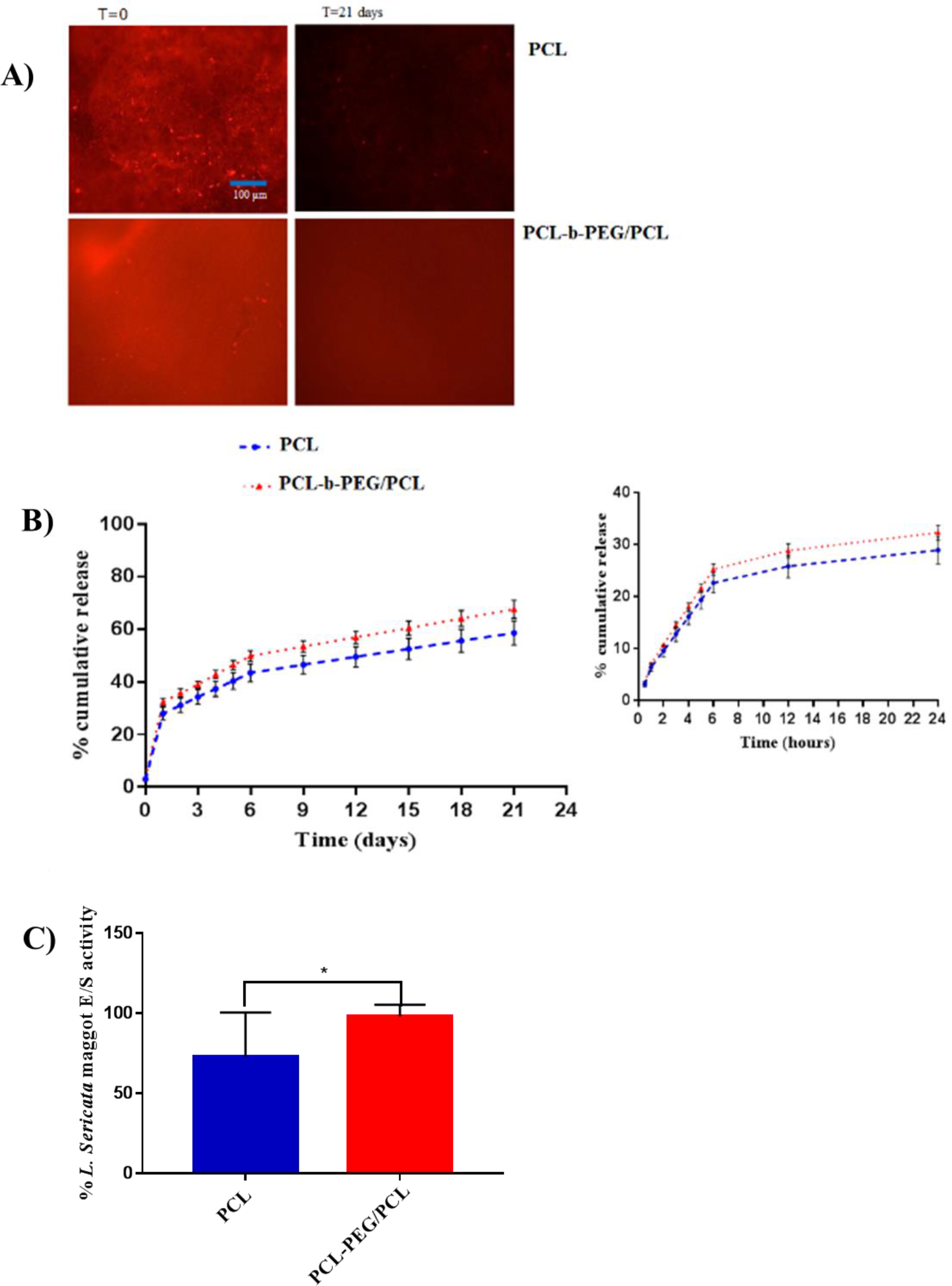
A) Fluorescent microscopy images of *L. sericata* Maggot E/S-rhodamine B conjugate protein absorbed on PCL and PCL-b-PEG/PCL scaffolds. Magnification 10x, scale bar 100 microns. B). *L. sericata* maggot E/S release profiles from PCL and PCL-b-PEG/PCL scaffolds in PBS pH 7.4 at 37°C. Experiments carried out in duplicated, and 3 independent experiments (means±SD). **C)** Relative *L. sericata* maggot E/S activity. Maggot E/S release pooled fraction from PCL and PCL-b-PEG/PCL scaffolds were analysed through protease assay. The data as *L. sericata* maggot E/S activity (%) normalised for protein content. Experiments carried out in triplicate, and three independent experiments (means±SD).

Figure 5-C shows the bioactivity of *L. sericata* maggot E/S after release from PCL and PCL-b-PEG/PCL scaffolds. Enzymatic activity of *L. sericata* maggot E/S released from PCL-b-PEG/PCL was (98.0 ± 2.6 %), which was significantly higher (P≤0.05) than the enzymatic activity from PCL scaffolds (72.9 ± 9.2%). The data was expressed as *L. sericata* maggot ES activity (%), with respect to the initial *L. sericata* maggot E/S activity which was assessed during the loading and after the released studies.

## Discussion

The design of bioactive scaffolds is an effective strategy to stimulate and promote tissue regeneration [21]. There is a high need to develop materials that are capable of supporting cellular growth whilst releasing therapeutic molecules to modulate cellular function and promote tissue regeneration [22].

Fibre diameter determines the pore size and porosity of electrospun scaffolds which play an important role as the structural support for cellular proliferation and infiltration [23, 24]. It has been reported that fibroblasts prefer scaffolds with a fibre diameter in the sub-micron scale as these scaffolds allow fibroblasts to penetrate into the fibres [25, 26] while electrospun fibre diameters on the nanoscale, reduces the average pore size and porosity [27, 28]. The scaffolds produced in this study are in line with the above values.

Blends of PEG-PCL polymers have been used as a strategy to modify the mechanical properties, wettability, biocompatibility, and solubility of these scaffolds in water and thus accelerate the degradability of the scaffolds [29, 30]. The mechanical properties of scaffolds designed for the treatment of skin lesions should match those of the skin itself [31]. For native skin, values of Young modulus are reported to be within the range of ~5-30 MPa [32]. The PCL and the PCL-b-PEG/PCL electrospun scaffolds presented here matched these mechanical properties. It was hypothesised that increasing the hydrophilicity of the scaffolds, by the incorporation of PEG, would also support cellular biocompatibility and increase the degradation rate of the material. The findings presented here are in agreement with studies conducted by Huan et. al 2004 [33], who compared the degradation of three different types scaffolds (fabricated by using a 3D dispensing rapid prototyping machine) made from PCL, PCL-b-PEG and PCL-PEG-PCL over 60 weeks. PCL-b-PEG scaffolds degraded faster than PCL due to PEG-rich species releasing soluble oligomers. In this study, no significant differences were found in terms of wettability between both materials (PCL and PCL-b-PEG/PCL scaffolds) however the measurement of wettability may have been affected by the roughness of the electrospun scaffolds. We found that the PCL-b-PEG/PCL significantly improved cellular infiltration and biocompatibility compared with PCL scaffolds which is in line with findings reported by others [29, 34]. These scaffolds had the same fibre diameter and therefore we attribute these finding to differences in chemical composition due to the moisture–rich environment that PEG creates around cells adherent to the scaffolds [35]. However, this is only true at certain concentrations of PEG; at higher concentrations we observed that the scaffolds were unable to support 3T3 proliferation. It is well accepted that certain concentrations of PEG decrease protein adsorption and therefore inhibits cell attachment and protein adsorption, due to an entropic steric repulsion effect [36].

This investigation also aimed to develop an electrospun scaffold loaded with therapeutic biomolecules in addition to supporting dermal fibroblast proliferation and infiltration into the material. For our intended application of wound healing, such a scaffold would provide structural stability to cell-growth and infiltration, as well the capacity of releasing active biomolecules. Biomolecules that have therapeutic properties, including antimicrobial, antioxidant, anti-inflammatory and mitogenic activities have been used for many years for wound healing [22]. There are a number of examples of active dressings based on electrospun scaffolds containing growth factors, antimicrobials and other drugs (i.e antioxidants) [37, 38].

We chose secretions from *L. sericata* maggot E/S that has been previously shown to enhance the migration of fibroblasts which are a pivotal in wound healing [39]. The incorporation of bioactive molecules secreted from *L. sericata* maggots E/S on the surface of the electrospun fibres of PCL-PEG/PCL fibres is a novel approach for skin wound healing and it is the first time that bioactive molecules from this organism has been included within electrospun scaffolds.

The data showed that *L. sericata* maggot E/S could be successfully loaded onto PCL-b-PEG/PCL scaffolds but this was not significantly different with respect to loading PCL scaffolds. In addition, the data illustrated that *L. sericata* maggot E/S was released in a sustainable manner from all scaffolds and to a greater extent from the PCL-b-PEG/PCL scaffold at all timepoints measured. At the end point (21 days) *L. sericata* maggot E/S cumulative release reached 67% of the original loading from PCL-b-PEG/PCL scaffolds and 58% from PCL with the majority released within the first 24 hours. In addition, the released *L. sericata* maggot E/S from PCL-PEG/PCL retained its biological activity. This concentration may be more than enough to enhance fibroblast migration. According to Horobin et al. (2006) 1 μg/mL of fresh *L. sericata* maggot E/S enhanced fibroblast migration *in vitro* [5]. In the present study, approximately 22 μg/mL of *L. sericata* maggot E/S were absorbed onto PCL-b-PEG/PCL scaffolds and from this 14 μg/mL of *L. sericata* maggot E/S were released with remained biological activity and have the potential to enhanced fibroblast migration.

The results presented here illustrated that PCL-b-PEG/PCL fibrous scaffolds supported mammalian cell adhesion, proliferation and cell infiltration within the fibres. In addition, PCL-b-PEG/PCL scaffolds were suitable for the loading, release and preservation of bioactivity of *L. sericata* maggot E/S. In the future, we aim to evaluate therapeutic effectiveness of the *L. sericata* maggot E/S released from such scaffolds on fibroblast, endothelial and keratinocyte proliferation though *in vitro* scratch assays. An *in vivo and ex vivo* model that allows the testing of the effect of *L. sericata* maggot E/S for a long period of time (including histological assessment) will be necessary to evaluate any long-term impact on wound healing.

## Conclusions

The development of a bioactive scaffold which combines a supportive structure for mammalian cell proliferation and infiltration in addition to being a drug release platform, is a promising approach for tissue engineering applications. Herein we showed that a biocompatible PCL-PEG/PCL fibrous scaffold supported skin fibroblasts and the release of *L. sericata* maggot E/S which includes enzymes previously shown to modulate fibroblast motogenesis and angiogenesis. In addition, PCL-PEG/PCL scaffolds used as a platform for drug delivery provides a stable environment to preserve the bioactivity and integrity of therapeutic molecules, including proteins.

The sustained release of serine proteinases from PCL-b-PEG/PCL fibre scaffolds is proposed for potential application in wound healing. The incorporation of bioactive molecules secreted from *L. sericata* maggot E/S on the surface of the electrospun fibres of PCL-PEG/PCL fibres is a novel approach for skin wound healing and it is the first time that bioactive molecules of *L. sericata* maggot have been included within electrospun scaffolds to produce a controlled release 3D scaffold.

## Acknowledgements

A Giacaman was supported by a grant from the ‘Comisión Nacional de Investigación Científica y Tecnológica de Chile’.

